# An automated two-choice social operant task for probing moment-to-moment changes in social satiety

**DOI:** 10.1101/2025.02.26.640376

**Authors:** Nancy R. Mack, Tomohito Minakuchi, Ken Igarza, Pauline Wonnenberg, Stefan N. Oline, Annegret L. Falkner

## Abstract

In naturalistic conditions, animals must routinely make ongoing self-motivated choices about whether to initiate interactions with same or opposite-sex partners and whether to re-initiate social contact when interactions have ended. However, it remains unclear what governs these choices, and whether they are motivated by drive states that exhibit signatures of moment-to-moment social satiety when these interactions have ended. Here, to explicitly test this at the behavioral level, we designed a novel fully-automated two-choice social operant paradigm where individuals can choose between same and opposite-sex social rewards and rewards are delivered for interaction with systematically varying durations. We trained cohorts of both sexes on this task and quantified the patterns of choices. We used choice latency as a metric to infer moment-to-moment satiety to test whether increased interaction duration leads to increased moment-to-moment satiety. We find that although both males and females have stable choice biases across sessions, with males showing consistent opposite sex biases, only males exhibit behavioral signatures of moment-to-moment social satiety and are sensitive to the duration of interaction. Using a simple normative model to capture patterns of social choice, we observe that behavior is better fit by a model that has a single evolving social drive and choice bias, rather than a model with multiple, independent drives for same and opposite-sex interactions.

Together, our data reveal behavioral signatures of social satiety and offer new insights into the underlying homeostatic and motivational drives that govern social choices.

---

The process of satisfying physiological needs such as fluid, salt, and other nutrients is often referred to as a “drive” state (Augustine, Lee, and Oka 2020). Individuals will adjust their moment-to-moment behavior in order to reduce a negative drive state (e.g. hunger or thirst) (Betley et al. 2015), and then pause consumption if they are sated, or fully satisfied. In models of homeostatic seeking behavior, which reflect the tendency of individuals to engage in behaviors to maintain a state of balance, elapsed time since the consumption is correlated with increased drive and the volume of the consumed reward is correlated with increased satiety and reduced drive (Keramati and Gutkin 2014). For example, during feeding behavior, individuals may pause consumption for longer, if a larger amount of food has been consumed (Rolls et al. 1998). In the social domain, specific manipulations across long timescales, including chronic isolation, appear to increase the likelihood of social interaction when animals are reunited, suggesting that absence of social contact may build up a “need” for social contact (Liu et al. 2023; Lee, Chen, and Tye 2021; Zelikowsky et al. 2018), which we refer to here as social drive. Additionally, sexual contact in males has been previously shown to generate biochemical signatures that promote prolonged abstinence from additional sexual contact, on the timescale of hours (Zhang et al. 2021). However, one major open question is whether brief social interactions can induce social satiety, which we define behaviorally as the latency between social choices, on a moment-to-moment timescale. Moreover, does the content or duration of a brief interaction predict moment-to-moment satiety?

A second major open question is whether social choice behavior is predicted by multiple independent parallel drives for competing needs, or by a single more generic “social” drive. For example, is the drive to interact with a same sex partner independent from the drive for social interaction with an opposite sex partner? On one hand, drives for same and opposite sex partners may be highly specific, since desired social outcomes during the interactions themselves may be divergent. For example, in males, drives for reproduction and aggression may be separable, and social choices towards opposite and same sex partners might be made to satisfy each of those drives independently. On the other hand, many drives have physiological or circuit-level dependencies. For example, cravings for salt and water are connected to each other (Lowell 2019). Given that social drives share many circuit-level features and hormonal influences, social drives may be similarly intertwined (Mong and Pfaff 2003; Rubin, Reinisch, and Haskett 1981). Answering this will require the use of normative models that make explicit predictions about how animals make self-motivated choices between different social targets that can account for a wide range of data.

To answer both of these questions we designed a novel, fully-automated two-choice social operant task, where mice learn that they can acquire brief interactions with either same or opposite sex partners. We trained cohorts of male and female mice, and by varying the duration of the social interaction in blocks, we tested whether differences in interaction duration induce moment-to-moment satiety by looking at the changes in choice and choice likelihood following those interactions. In addition, we examined whether a pair of simple normative models that assume either a single social drive or multiple independent drives better predicts behavior. We found that choice could be well-described with only a few key features, including choice bias and interaction probability. Surprisingly, we found that males are highly sensitive to satiety manipulation, exhibiting behavioral signatures of moment-to-moment satiety, while females are not. However in both cases, data was better fit by a model with a single social drive rather than multiple independent drives. These findings provide critical insight into the underlying homeostatic and motivational states that drive social choice.

## Male and female mice learn the 2-choice SOAR task

We created a fully automated two-choice social operant task where mice can nose poke for access to a male or female conspecific for a specified duration of time (Supplementary Video 1). The operant chamber is composed of two sides, each outfitted with a “social port” and a “null port.” A nose poke into the social port resulted in automated delivery of a conspecific from above into the chamber for a fixed duration of time, after which the reward animal was automatically retracted back to a position outside the chamber (Fig 1A-B). Social rewards are cradled in a body-length jacket that provides support and restricts movement of the social target, yet allows the reward animal to support its own weight, and to be investigated and interacted with by the experimental animal. We refer to this interval of potential social interaction as the “reward down time,” and the “interpoke interval (IPI)” as the time between the interaction end and the next poke (Figure 1B), akin to an intertrial interval. Poking the null port resulted in no change in task variables and is used to infer learning. The task is entirely self-paced by the subject animal, who freely makes choices throughout the duration of the session with no external motivational incentives (i.e. food or water). We trained cohorts of male and female CD1 mice on this task by first training them on a single-choice version of the task. To do this, we restricted access to a single side of the operant chamber with a removable wall. Mice were first trained on a single side for 3-6 days until task criteria were reached, and then were trained on the opposite side for a similar duration (Fig 1C). Training for the female or male-side first was counterbalanced across groups. We used pose tracking (SLEAP) to track the position of the subject and reward animals in the operant chamber using videos recorded during each session using top and side cameras ^12^.

**Figure 1:**
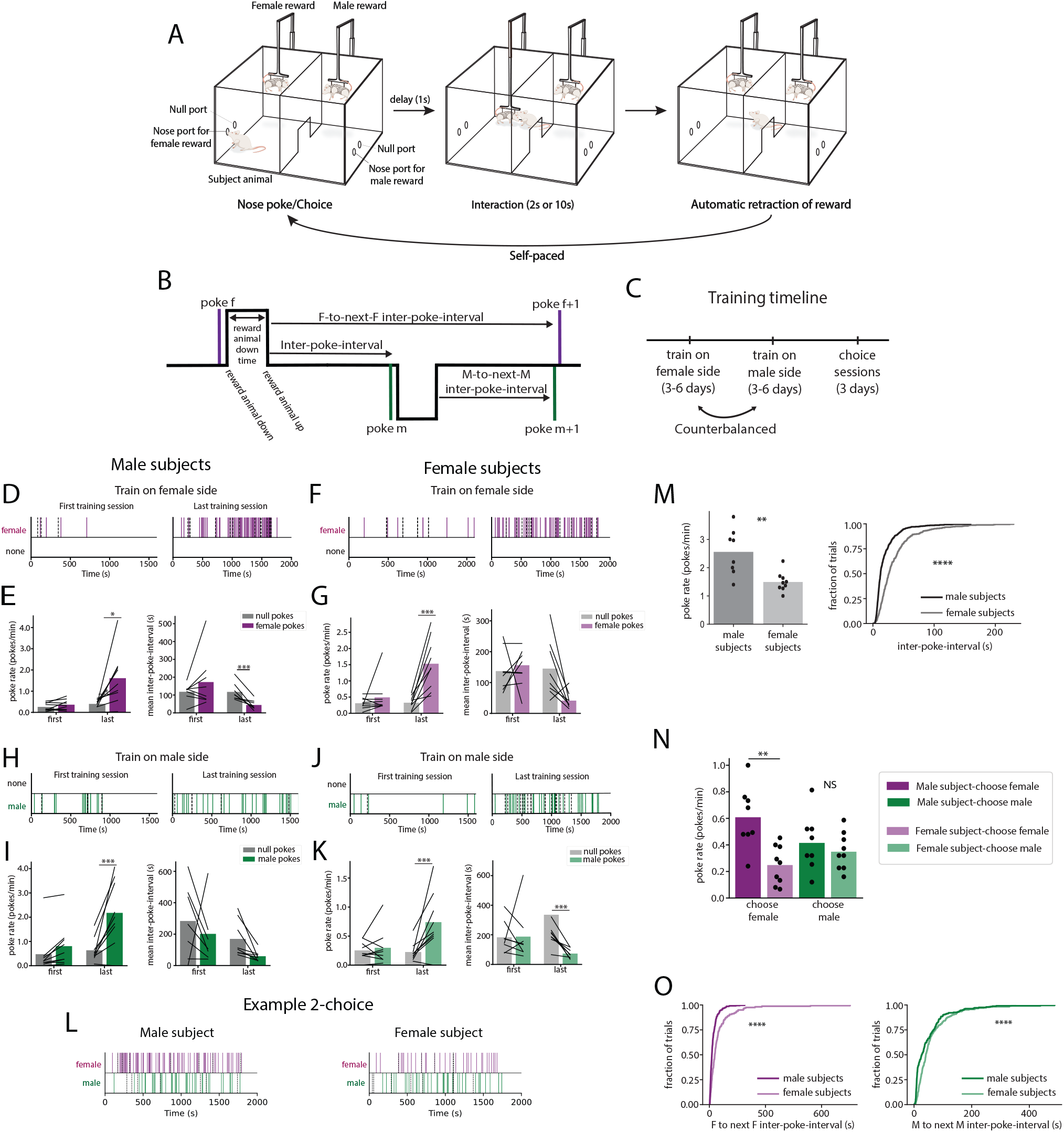
Males and females are trained and tested using the SOAR task. **(A,B)** Schematic of the 2-choice SOAR task. **(C)** Training timeline**, (D,E)** Male subjects (*n* = 9) training to poke for female rewards. (**D)** Representative raster plots showing nose-pokes of social (purple lines) and null pokes (dashed lines) over time on the first (left) and last (right) days of training. **(E)** Left: Change in poke rate on the social (purple) and null port (gray) for male subjects during training for female rewards (two-way mixed ANOVA with FDR correction for multiple comparison; social vs null on first day (*p* = 0.213) or last day (*p* = 0.0016)). Right: Change in mean latency between social pokes or null pokes on the first and last day of training for female rewards in male subjects (two-way mixed ANOVA with FDR post hoc test for multiple comparison; social vs null on first day (*p* = 0.453) or last day (*p* = 0.0456). **(F,G)** Female subjects (*n* = 9) training to poke for female rewards. (**F)** Representative raster plot from female subject depicting nose pokes across training as in **D. (G)** Left: Change in poke rate on the social (purple) and null port (gray) for female subjects during training for female rewards (two-way mixed ANOVA with FDR post hoc test for multiple comparison; social vs null on first day (*p* = 0.345) or last day (*p* = 0.0023). Right: Change in mean latency between social pokes or null pokes on the first and last day of training for female rewards in male subjects (two-way mixed ANOVA with FDR post hoc test for multiple comparison; social vs null on first day (*p* = 0.836) or last day (*p* = 0.011). **(H, I)** Male subjects training for male rewards. **(H)** Representative raster plots as in **D** depicting male pokes in green and null pokes in gray. **(I)** Left: Change in poke rate on the social and null port for male subjects training for male rewards. Right: Change in mean latency between social pokes and null pokes across training. **(J, K)** Female subjects training for male rewards. **(J)** Representative raster plot from a female subject depicting nose pokes across training for male rewards. **(K)** Left: Change in poke rates across training (two-way mixed ANOVA with FDR correction for multiple comparisons; social vs. null on first day (*p* = 0.589) or last day (*p* = 0.016). Right: Change in mean latency between social or null pokes across training. (**L)** Example raster plots during a two-choice SOAR session. (**M)** Males show greater total pokes/min during a two-choice SOAR session compared to female mice (independent samples t-test, *p* = 0.0027). Right: Cumulative distribution of interpoke-intervals between all pokes is shifted leftward in female subjects compared to males (Kolomogorov Smirnov test, *p* = 5.5e-30). (**N)** Males have a higher poke rate for female choices compared to female subjects (*p* = 0.0058), but equal poke rates for male choices (*p* = 0.943, two-way mixed ANOVA, pairwise comparison with FDR correction). **(O)** Cumulative distribution for interpoke-intervals is shifted leftwards in female subjects compared to males subjects for female choices (left, purple, Kolomogorov Smirnov test, *p* = 1.85e-11) and male choices (right, green, Kolomogorov Smirnov test, *p* = 2.04e-08). Data in bar graphs are presented as mean values. * p < 0.05, ** p < 0.01, *** p < 0.001.

We found that both male and female subjects readily learned this task (Fig. 1D-K). On the first day of training for same or opposite sex rewards, poke rates and interpoke intervals for the social and null port were similar in both male and female subject mice (Fig 1D-K, left). However, on the last day of training, both sexes showed significantly higher poke rates and lower interpoke intervals on the social port compared to the null port (Fig 1D-K, right). Importantly, in a single-choice version of this task, we have previously shown that mice will not nosepoke for the arm to move in the absence of a stimulus mouse being present, or for a novel object (Falkner et al. 2016; Minakuchi et al. 2024). Together, these data indicate that both sexes learned to discriminate between the social and null port for access to a female or a male conspecific.

## Sex differences in choice behavior during the two-choice SOAR task

If individuals reached training criteria on each side of the single-choice task, they were tested on the “full” 2-choice version of the task for 3 successive days where they had access to both same and opposite sex rewards (Fig. 1L). We describe behavior in this task within each session by quantifying a few key features: 1) Total poke rate across all choices, 2) Poke rate for specific choice (poke rate male and poke rate female, 3) Interpoke interval (the length of time from the reward retraction to a specified choice type), 4) Interaction probability (likelihood that interaction time contained a direct social interaction), and 5) Choice bias (defined as the proportion of females choices out of the total, where 0.5 is an equal number of male and female choices in the session). We compared behavior across three sessions of the two-choice SOAR task and observed that social poke rates increased across days for both sexes while null poke rates were consistently low (Fig S1). To evaluate potential sex differences, we compared performance on the final 2-choice SOAR session.

In the 2-choice SOAR task, we observed that male subjects have significantly higher total poke rates and shorter interpoke-intervals compared to female mice (Fig 1M). By breaking down poke rates and interpoke-intervals by choice type, we observed that male subjects have higher poke rates for female rewards, while both sexes show similar poke rates for male rewards (Fig 1N). However, males exhibit shorter interpoke-intervals between the same choice type (F-F, M-M) for both choice types compared to female subjects (Fig 1O). Overall, these results show that males choose female rewards more frequently in the two-choice SOAR task compared to female subjects, and that regardless of choice type, male subjects show shorter intervals between choices compared to female subjects.

Next to examine whether choice bias is a stable property of an individual, we compared choice bias across the 3 days of the 2-choice task (Fig 2A-C). Surprisingly, we found that both sexes exhibit relatively stable choice bias across days, with similar variance observed between male and female subjects (Fig 2B). On average, we found that males have a higher choice bias (mean = 0.61 ± 0.048) than female subjects (M=0.42 ± 0.060) and male are biased towards female preference across the population, while females are not biased (Fig 2C). Though males were typically female-biased, females were sometimes stably male-biased, and sometimes stably female-biased, suggesting that this bias is a stable property of an individual.

**Figure 2:**
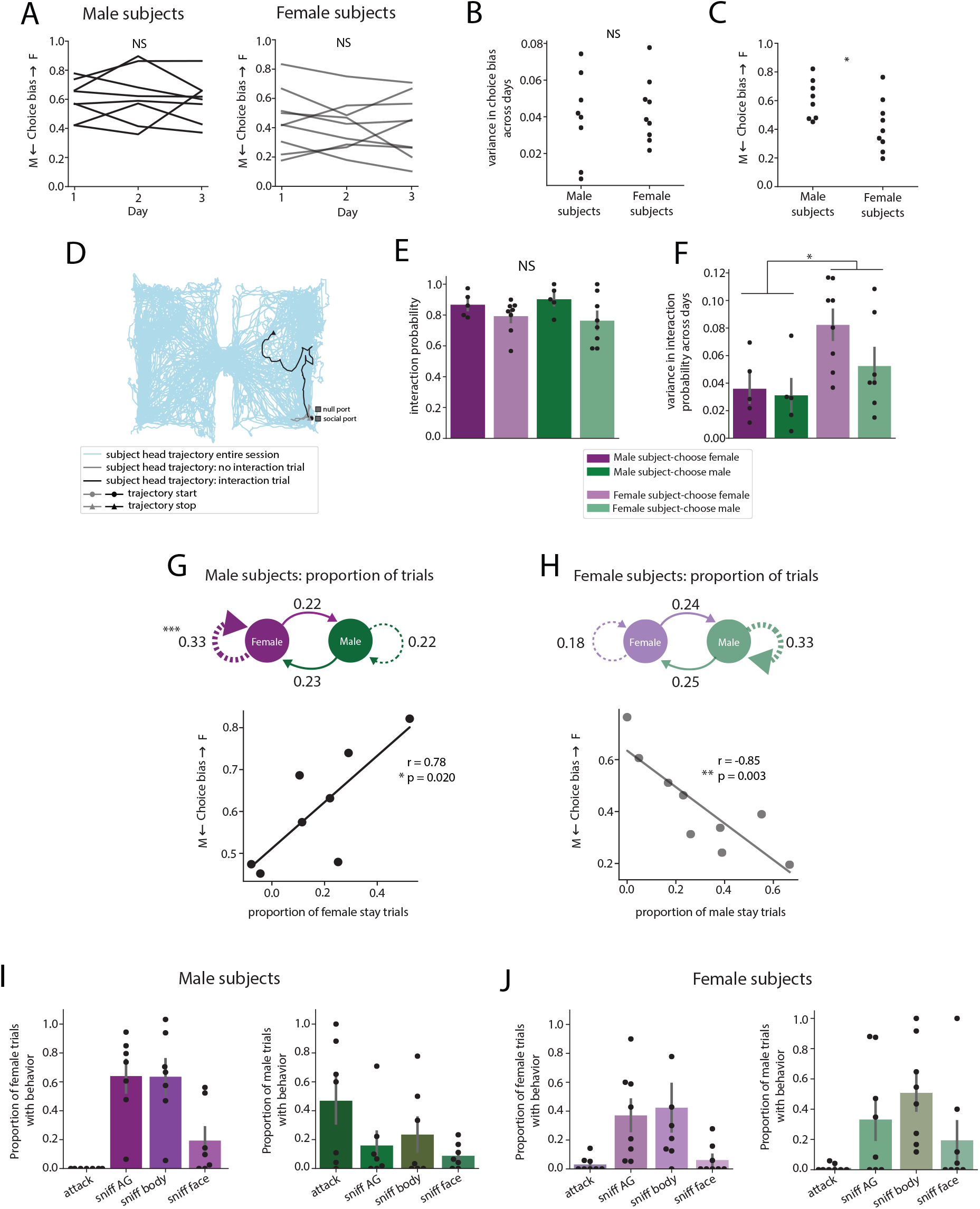
Behavior in 2-Choice SOAR task can be described by choice bias, choice probability, and interaction probability. **(A)** Choice bias (# female pokes/#total pokes) is stable across the three two-choice SOAR sessions for male subjects (left, one-way repeated measures ANOVA, *p* = 0.893) and female subjects (right, one-way repeated measures ANOVA, *p* = 0.652). **(B)** The variance in choice bias across days is equal between male and female subjects, (independent samples t-test, *p* = 0.748). **(C)** Males have a greater choice bias (skewed towards female) relative to female subjects, *t* = 2.32, *p* = 0.0346. **(D)** Plot showing tracking of subject animal’s head in 2-choice operant chamber (blue) throughout the duration of the task, with an overlaid example of the subject’s head during a male trial with interaction or no interaction. **(E)** There are no sex differences in interaction probability following female choices (purple) or male choices (green), two-way mixed ANOVA, *F*_(1,11)_ = 0.673, *p* = 0.429. **(F)** Females show a greater variance in interaction probability across days for male and female choices relative to male subjects. **(G)** Top: Transition plot between trials with interaction. Males show the greatest proportion of female-stay trials. (Chi-square test, *p* = 0.00024). Bottom: There is a significant, positive correlation between an individual’s mean choice bias and average proportion of female-stay trials, pearson’s r = 0.78, p = 0.020. **(H)** Top: Same as D but for female subjects, who show the greatest proportion of male-stay trials (Chi-square test, *p* = 0.1053). Bottom: Female subjects show a negative correlation between mean choice bias and average proportion of male-stay trials, pearson’s r = −0.85, *p* = 0.003. **(I)** Proportion of female trials (left, purple) or male trials (right, green) containing attack, sniff anogenital (AG), sniff body, or sniff face behavior in male subjects. **(J)** Same as **(I)** but for female subjects. Bar graphs are presented as mean values ± s.e.m. * p < 0.05, ** p < 0.01, *** p < 0.001. NS, nonsignificant.

Next, since individuals can poke, but not actually interact with the social reward, we quantified whether same or opposite sex choices resulted in interactions during the reward-down-time by using pose to track the position of the subject’s head following each choice (Fig 2D). While interactions did not occur on every trial, both sexes showed a similarly high probability of interacting with both male and female reward animals (Fig 2E). To compare the stability of interaction probability across days, we found that female subjects exhibited higher variance in the probability of interaction for both male and female rewards relative to male subjects (Fig 2F). However, this variability cannot be explained by differences in the estrous state of the female subjects (Fig S1E). We next manually annotated the behaviors displayed by the subject mice during male and female trials with interaction (Fig 2I,J). We observed typical interaction behaviors, including sniffing of the anogenital region, body, or face of the social reward animals in both male (Fig 2I) and female subjects (Fig 2J). Several male subjects showed high probability of attack following male choices, but never for female choices (Fig 2I). In female subjects, attack behavior following male or female choices was extremely rare. These findings show that interactions in this operant task are similar to what would be expected or observed in more naturalistic social contexts.

Requesting the same social choice repeatedly may indicate a lack of satiety for that social reward following the end of the interaction. To quantify this across male and females, we compared the relative proportion of trials with interaction where individuals made the same choice (“stay” trials) compared to trials where they changed their social request (“switch” trials). Male subjects show the highest proportion of “female stay” trials, female choices that are followed by another female choice (Fig 2G top). As expected, we found that the mean proportion of female stay trials across days showed a positive correlation with individual choice bias, such that animals with the largest female preference displayed the greatest proportion of female-stay trials (Fig 2G, bottom). In female subjects, we observed the opposite trend, such that females had greater than expected proportions of male-stay trials that, on a per animal basis, showed a negative correlation with individual choice bias (Fig 2H). Altogether, these results demonstrate that both sexes show a “stay bias:” for the opposite sex.

## Social satiety manipulations influence choice bias and interaction probability

The fully automated nature of the two-choice SOAR task allows experimenters to systematically vary the duration of potential social interaction (the reward-down time) separately for each choice type on a trial-to-trial or block basis. We used this parameter to probe whether longer reward-down times induced changes to moment-to-moment social satiety by testing the hypothesis that longer reward-down times increase satiety. We quantified moment-to-moment satiety by assessing changes in choice probability and the latency to the next choice (IPI). We used a block design across 3 successive days to compare the differences between equal, male, and female satiety manipulations across both sexes (Fig 3A). During the “equal” block, both male or female choices resulted in short reward-down times (2 seconds, f-short, m-short). During the “female satiety” block, female-directed choices resulted in long reward-down time (10s, f-long) while male choices resulted in short reward-down times (2s, m-short). Conversely, the “male satiety” block consisted of long male reward-down times (10s, m-long) and short female reward-down times (2s, f-short). We categorized trial types by their current choice followed by their successive choice (Fig. 3B, e.g. F-F represents a female-directed choice followed by another female-directed choice). Latencies for specific trial types could also be classified without respect to the next choice (F-to-any, M-to-any).

**Figure 3:**
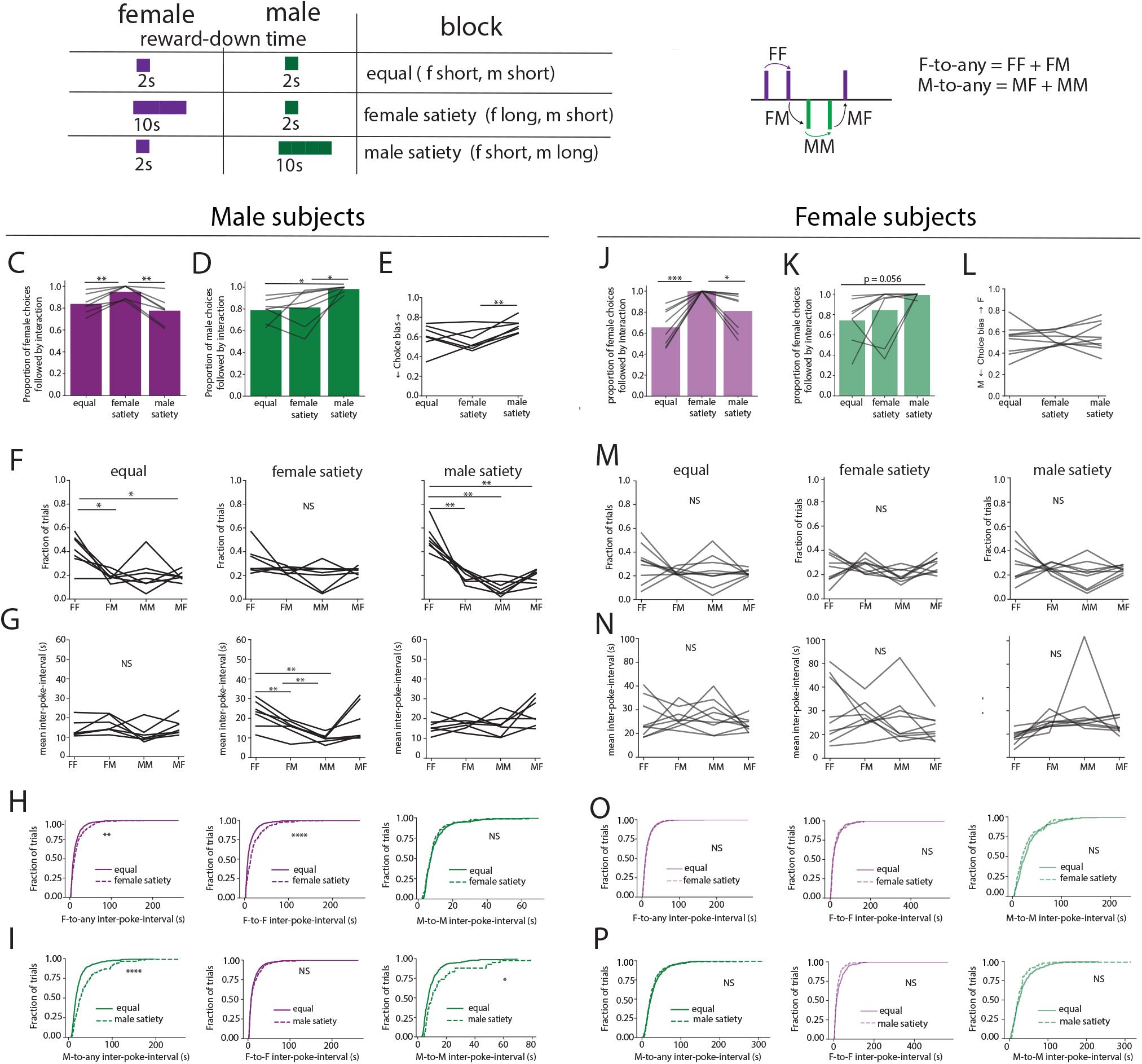
Males but not females are sensitive to changes in moment-to-moment social satiety. **(A)** Schematic of different experimental conditions used to probe social satiety. In equal reward sessions, the interaction time for male or female rewards were both set to 2 seconds. In Female long sessions, interaction time with the female reward was set to 10 seconds while interaction with the male reward was set to 2 seconds. In male long sessions, male interaction time was set to 10 seconds while female interaction time was set to 2s. **(B)** Example behavior raster schematizing different trial types within a session, defined by the current poke and the next poke. FF: female-to-female, female pokes followed by another female poke, also considered a “female stay” trial. FM: female-to-male, male poke followed by a male poke, also considered “female switch” trial. MM: male-to male, male poke followed by another male poke, or “male stay” trial. MF: male-to-female, male poke followed by a female poke, or “male switch”. C-I, Male subjects (*n* = 8) tested during equal, female satiety, or male satiety blocks. **(C)** The proportion of female choices followed by social interaction is the highest during female satiety, (one-way repeated measures ANOVA, *p* = 0.00036. Pairwise comparison with FDR correction, equal vs. female satiety: *p* = 0.0056; female satiety vs. male satiety: *p* = 0.0056). **(D)** The proportion of male choices followed by social interaction is highest during the male satiety block (one-way repeated measures ANOVA, *p* = 0.0098. Pairwise comparison with FDR correction, equal vs. male satiety: *p* = 0.0133; female satiety vs. male satiety: *p* = 0.0486). **(E)** Choice bias in male mice is most skewed towards female preference during the male satiety block (one-way repeated measures ANOVA, *p* = 0.0194. Pairwise comparison with FDR, female satiety vs. male satiety: *p* = 0.0182). **(F)** Proportion of trial types during the equal (*left*), female satiety (*middle*), or male satiety (right) blocks. During the equal block (*left*), male mice show the greatest proportion of FF trials (one-way repeated measures ANOVA, *p* = 0.0056. Pairwise comparison with FDR correction, FF vs. FM: *p* = 0.025; FF vs. MF: *p* = 0.025), but during the female satiety block (middle), the proportion of different trial types is equal (*p* = 0.131). During the male satiety block (*right*), males show the highest proportion of FF trials. (One-way repeated measures ANOVA, *p* = 3.056e-07. Pairwise comparison with FDR correction, FF vs. FM: *p* = 0.0038; FF vs. MF: *p* = 0.00167; FF vs. MM: *p* = 0.0037). **(G)** Mean interpoke-intervals (IPIs) for different trial types during the equal (*left*), female satiety (*middle*), or male satiety (right) blocks. There are no differences in mean poke latencies across different trial types during the equal block (*left,* one-way repeated measures ANOVA, *p* = 0.0933). During female satiety, the mean interpoke-interval is greater for FF and FM trial types (*middle,* one-way repeated measures ANOVA, *p* = 0.0015. Pairwise comparison with FDR; FF vs. FM:, *p* = 0.0097; FF vs. MF: *p* = 0.00737; FM vs MF: *p* = 0.0097. During male satiety, there is no difference in mean interpoke-intervals across trial types (right, one-way repeated measures ANOVA, *p* = 0.140). **(H)** Distribution of IPIs for different choice sequences between equal blocks (solid line) or female satiety block (dashed line). For female choices, there are longer IPIs to the next trial (left, Kolomogorov Smirnov test, *p* = 0.0063) or the next female trial (middle, Kolomogorov Smirnov test, *p* = 5.67e-07) during the female satiety. For male choices, IPIs to the next male choice is similar during female satiety block relative to equal blocks (right, Kolomogorov Smirnov test, *p =* 0.644). **(I)** Distribution of IPIs for different choice sequences between equal blocks (solid line) or male satiety block (dashed line). During the male satiety block, there are longer IPIs from male choices to the next choice (*left*, Kolomogorov Smirnov test, *p =* 9.55e-08) or the next male choice (*right*, Kolomogorov Smirnov test, *p* = 0.0257) relative to equal blocks. For female choices during male satiety, IPIs to the next female choice are similar to that during equal blocks (*middle,* Kolomogorov Smirnov test, *p* = 0.546). **J-O,** Female subjects (*n* = 9) tested during equal, female satiety, or male satiety blocks. **(J)** The proportion of female choices followed by social interaction is highest during female satiety relative to the equal block or male satiety block (one-way repeated measures ANOVA, *p* = 0.002. Pairwise comparison with FDR correction, equal vs. female satiety: *p* = 0.00225; female satiety vs. male satiety: *p* = 0.036). **(K)** There was a trend towards increased trials with interaction following male choices during the male satiety block that did not reach statistical significance (*p* = 0.0561). **(L)** Female subjects do not show a significant change in choice bias across the different block sessions (one-way repeated measures ANOVA, *p* = 0.878). **(M)** There are no significant differences in proportion of choices or **(N)** mean interpoke-interval across different trial types during equal (*left*), female satiety (*middle*) or male satiety (*right*) blocks. **(O)** There is no difference in the distribution of IPIs for female choices to the next choice (*left*) or next female choice (*middle*), or from male choices to the next male choice (*right*) during equal blocks relative to the female satiety block. **(P)** The distribution of IPIs for male choices to the next choice (*left*) or next male choice (*middle*), or from female choices to the next female choice (*right*) are similar during equal blocks relative to the male satiety block. NS, nonsignificant.

First to verify that longer reward-down times would lead to increased social interaction, we compared the interaction probability for each reward type across the three satiety blocks. We observed that in male subjects, longer reward-down in the male satiety block was associated with increased interaction probability following male choices when compared to the shorter reward-down time for males, and longer reward-down time in the female satiety block was associated with increased interaction probability relative to the shorter reward-down times for females (Fig 3C-D). A similar trend was observed in female mice (Fig 3J-K). Next, we examined whether satiety manipulations would change choice bias, by comparing choice bias in male and female satiety blocks. In male subjects, we observed a significant change in choice bias across the different blocks (Fig. 3E), with the lowest female preference observed during the female satiety block (M=0.56 ± 0.038 mean+/-SEM) relative to the equal block (M=0.60 ± 0.046) and male satiety block (M= 0.65 ± 0.025). In female mice, we did not find a shift in choice bias across the different block sessions (Fig 3L). These data suggest that while both sexes “take advantage” of the increased potential for social interaction, only males shift their choice bias in the opposite direction.

## Males exhibit behavioral signatures of social satiety while females do not

We next tested the specific prediction that long reward-down times induce greater moment-to-moment satiety relative to short reward-down times. This predicts that long reward-down times will result in increased interpoke interval for the following choice and a reduced likelihood of a “stay” trial for the previous choice (M-M or F-F). We found that during equal blocks, male subjects showed the greatest proportion of female-stay (F-F) trials relative to all other choices, and this preference was abolished during female satiety blocks exclusively (Fig 3F). Indeed, the proportion of female stay trials relative to all female choices was significantly reduced during female satiety blocks compared to equal and male satiety blocks. We also quantified the latency to initiate another choice following the end of social interaction (IPI) across choice types across different block sessions. During equal blocks, male mice showed similar mean IPIs across all trial types (Fig 3G, left). However, during the female satiety block, male subjects had greater mean IPIs for female stay and female switch trials relative to male-stay and male switch trials (Fig 3G, middle).

One prediction of moment-to-moment social satiety is that we should not observe relatively increased latencies following trial types that are not manipulated (short reward-down times, when the opposite choice is long reward-down). For example, if individuals are sensitive to satiety, long reward-down times for female choices should increase subsequent interpoke-intervals, but interpoke-intervals following male choice trials (that have a short reward-down time) should exhibit no change relative to equal blocks, since they are still expected to induce low levels of satiety. Indeed, when we compare the IPIs across satiety blocks, we observe increased IPIs following female trials relative to similar choices in the equal block (Fig 3H, left, middle), but the IPIs following male choices are not affected (Fig 3H, right). Similarly, in male satiety blocks, we observe that IPIs to the next male trial (Fig 3I, right) or to any trial type (Fig 3I, left) are increased, yet IPIs following female choices are not affected (Fig. 3I, middle). These data clearly indicate that males are sensitive to the satiety manipulation, and exhibit concomitant and specific behavioral changes (delayed reinitiation of social choice) that reflect this moment-to-moment satiety.

In stark contrast to male subjects, in female subjects, we observe almost no significant changes to behavior in choice proportion, choice-type IPI, or cross block IPI (Fig 3L-O) following satiety manipulations for same or opposite sex choices. These data suggest females may be insensitive to these small moment-to-moment changes in interaction-induced satiety.

## Modeling behavioral choice with 1-drive and 2-drives

A major open question is whether choice behavior is governed by multiple parallel drives for same and opposite sex rewards, or can be explained by a model with a single social drive state. We developed a pair of simple normative models (Fig. 4A-B) to simulate behavioral choices governed by each of these two hypotheses, and compared the simulated choice data from these models to data from both sexes across satiety manipulations.

**Figure 4:**
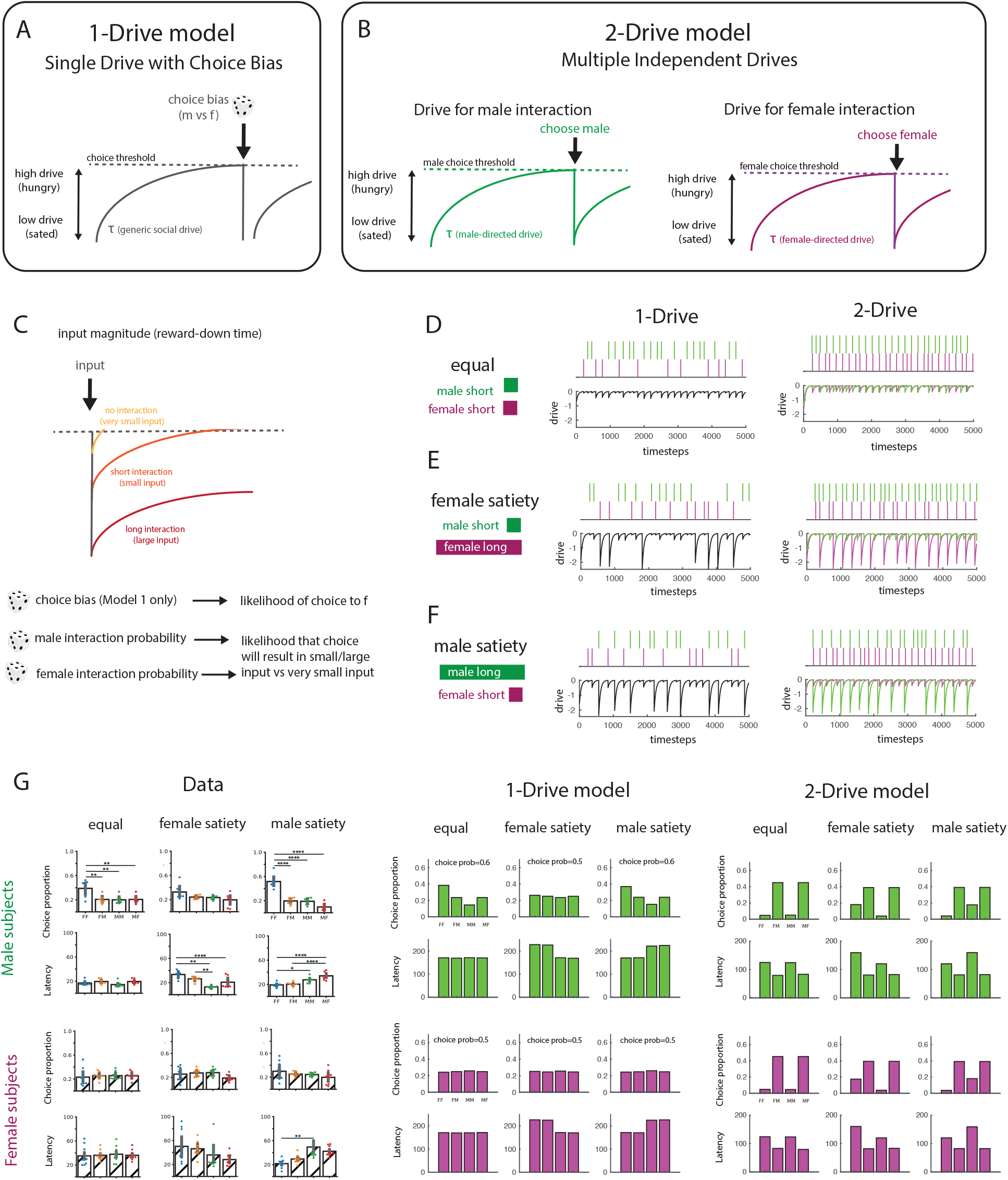
A normative single drive model with choice bias outperforms a multi drive model in predicting behavior across satiety states. **(A-B**) Descriptions of the 1-Drive (**A**) and 2-Drive models (**B**). In both models, drive is modeled as a state that evolves across time where input reduces this drive away from a threshold. In both models, a social choice is initiated when drive crosses a threshold. In the 1-Drive model **(A**), this is represented as a single drive variable, and choices between males and females are determined probabilistically by a fixed choice bias (**A**). In the 2-Drive model (**B**) drives for male and female choices evolve independently from each other, with separable thresholds for decision and decay constants. (**C**) In both the 1-Drive and 2-Drive models, input can vary in magnitude, which represents interaction duration, a proxy for satiety, and is defined by variable parameters that represent the likelihood of interaction with a female or a male. (**D-E**) Example simulations of the 1-Drive (**D**) and 2-Drive (**E**) models across the three satiety experiments: equal rewards with short interaction durations for both male and female trials (**D-E**, left), female satiety, with female choice associated with a large input (**D-E**, middle), and male satiety with male choice associated with a large input (**D-E**, right). For each simulation, choice rasters to male (green) and female (purple) choices are shown on the top plots, while the drive(s) are shown on the bottom plots. (**F-G**) Model simulations for 1-Drive and 2-Drive models relative to data for both choice proportions (top rows) and choice latencies (bottom rows). Data from males (**F**, left) and females (**G**, left) are better captured by the 1-Drive model, which recapitulates the key aspects of the data across satiety states The 1-Drive model captures the bias towards opposite sex stay trials, and the longer latencies associated with the sated choice, while the 2-Drive model predicts higher proportions of switch trials with short latencies.

The models were designed to predict choice using as few parameters as possible. Both models (1-Drive and 2-Drive models, Fig 4A-B) represent drive state as a state as a vector that decays towards 0 following an input, with more negative values indicating greater satiety (low drive) and values close to 0 indicating high drive (a “social hunger”, unsated state). In both models, when the drive value crosses a specific choice threshold, a choice is triggered. In the 1-Drive model, which assumes a single drive value, this choice is determined randomly (male choice or female choice), but is influenced by a stable choice bias term, set between 0 and 1 for a given simulation. For example, if the choice bias term is 0.5, the choice is equally likely to be male or female, but when the choice bias term is 0.6, the choice made at the time of threshold crossing is more likely to be female. Alternatively, in the 2-Drive model, which assumes that choices for males and females are driven by independent, parallel drives, these values evolve simultaneously with no choice bias term.

Input added to each model after each choice represents the social interaction itself and moves the continuous drive vector in the direction of satiety (Fig 4C). Small inputs represent short interaction times (i.e. the short reward-down times in the experiment) while large inputs represent long interaction times (i.e. long reward-down times). Since subjects do not socially interact in every trial (see Fig. 2G), we add an interaction probability term for each choice type in both models (male interaction probability and female interaction probability). Interaction probabilities were fixed to approximate real data, and were simulated using a random draw at each choice point. Input for no interaction trials was very small, but non-zero. We used both models to simulate the evolving drive and choices for the three satiety blocks, equal (Fig 4D), female satiety (Fig 4E), and male satiety (Fig. 4F). For each block type, we simulated the choice rasters single drive state for the 1-Drive model, and the choice rasters and parallel drives for the 2-Drive model.

We compared the output of each model for each satiety block to the data for both sexes, using parameter settings for choice bias and interaction probability that match the observed data means. For each simulated block, we quantified the model performance by looking at the choice proportion across each trial type, defined by the current choice and successive choice (F-F, F-M, M-M, M-F) and by the response latency/interpoke interval for each choice type (Fig 4G). We found that the 1-Drive model, but not the 2-Drive model captured several important features of the data. First, the 1-Drive model accurately predicts that males will have significantly more F-F stay trials than the other trial types, particularly when choice bias is female biased. In contrast, the 2-Drive model predicts that animals will have more M-F and F-M switch trial types, because their parallel drives will motivate them to alternate choice types. Second, the 1-Drive model predicts that latencies across trial types will be equivalent in the equal block, and will be longer following the stated trial type in the unequal blocks. For example, latencies will be longer for F-F and F-M trials during female satiety blocks, and latencies will be longer for M-M and M-F in male satiety blocks. Importantly, this feature is true even when choice bias is equal (0.5). In contrast, the 2-Drive model predicts that the “stay” type trial transition (F-F and M-M) will always be longer, regardless of the block type (Fig 4G), which is not supported by the data.

## Model Fits

Some parameters for initial simulations (including choice bias and interaction probability) were chosen as a match to data. However, with this strategy we can not rule out that some other parameter choice for the 2-Drive model would be a better fit to the data. To test whether some alternative combination of parameters for the 2-Drive model (choice threshold values, male tau and female tau decay terms, male interaction probability, female interaction probability), would accurately describe behavioral data in the 2-choice SOAR task, we simulated the runs of the model to iterate through combinations of these parameters for each satiety manipulation. All models were compared to the population means for the choice probability and choice latency from the data (Fig. 5A) using linear regression.

**Figure 5:**
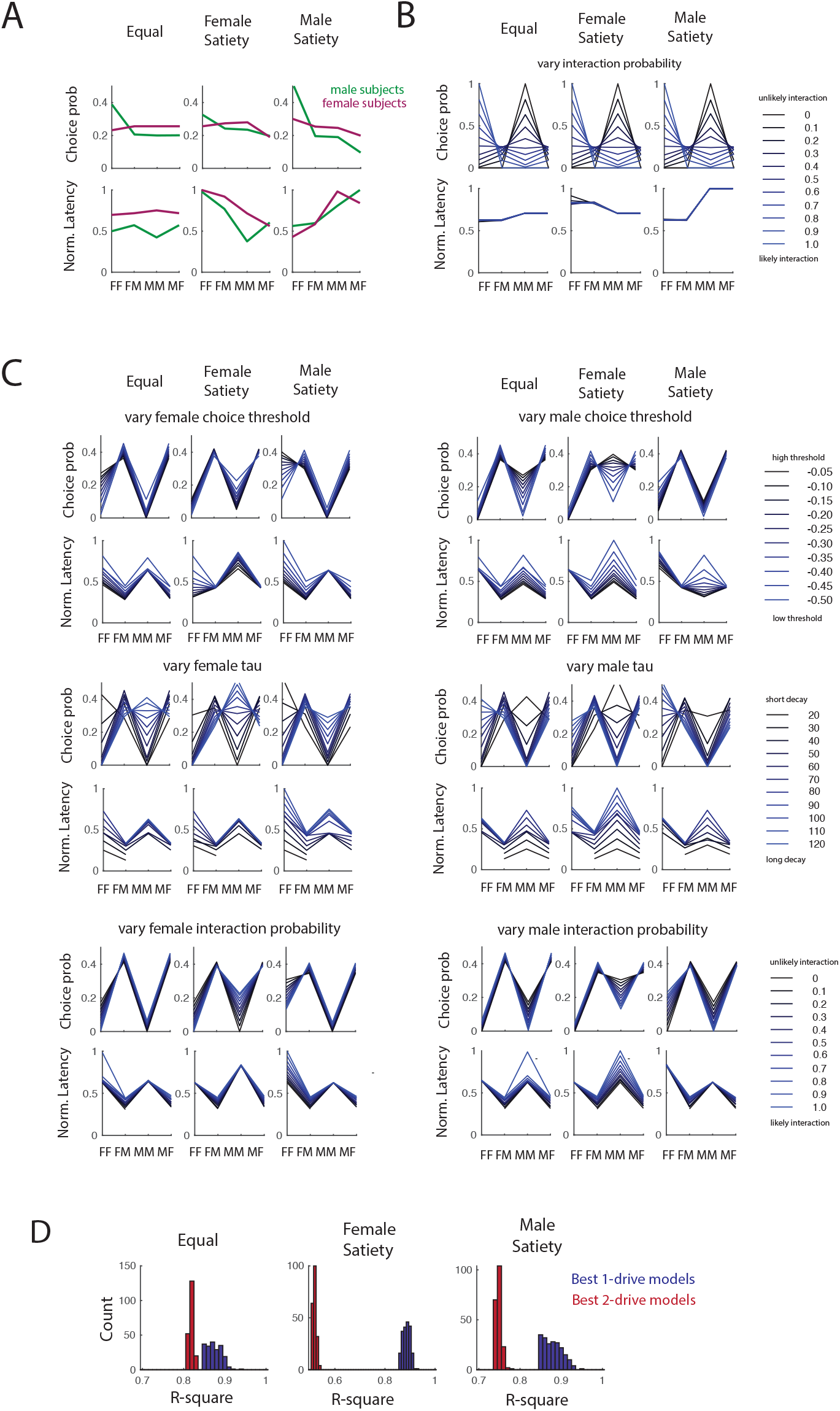
Varying parameters in the 2-Drive model does not describe behavior better than the 1-Drive model. (**A**) Mean choice probability (top) and latency (bottom) for 4 choice types (FF, FM, MM, MF) for males (green) and female (purple) for each satiety manipulation. (**B**) Simulations of 1-drive model across changes in the interaction probability. (**C**) Stimulations of the 2-drive model across changes in the choice thresholds, decay constants (tau), and interaction probabilities for male and females. (**D**) R-squared values of top 200 best fit models for the 1-Drive model (blue) and 2-Drive model (red) for each satiety manipulations.

For the 1-Drive model, we iteratively simulated across variations in interaction probability, and choice bias (Fig. 5B). This varied how likely a choice leads to either a small or large input compared to the very small input, the proxy for no interaction. For the 2-drive model, we iterated simulations across changes in the choice threshold for each drive, the tau for each drive, and the interaction probability for each choice type (Fig. 5C). Finally, we compared the outputs of these simulations to the data by fitting the simulated data to the choice probability and latency vectors. Using the fits of this linear model (R^2^), we compared the top 200 best fit 1-Drive and 2-Drive models (Fig. 5D). We found that across all satiety manipulations best fit 1-Drive models were better than best-fit 2-Drive models (Fig. 5D). This indicates that no combination of parameters in the 2-Drive model can out-perform the 1-Drive model at describing the behavioral data.

## DISCUSSION

Here we describe our use of a two-choice social operant task to probe moment-to-moment changes in self-motivated choices for same and opposite-sex social targets. Using this paradigm, we find that both male and female mice repeatedly self-initiate trials for interaction with both targets and show the highest frequency of stay trials for the opposite-sex reward. We observed that modeling social drive as a single drive state rather than with multiple, parallel drives recapitulated the observed greater frequency of same-type bouts, particularly in male mice. Furthermore, we found that males exhibited behavioral changes reminiscent of moment-to-moment changes in social satiety with key features of the dataset captured by the single drive model.

Similar to other single-choice social operant paradigms^13–19^, our data shows that mice will work to gain access to social targets of the same and opposite-sex, indicating that mice indeed find both types of interaction rewarding. Our observation that males have higher poke rates specifically for female rewards relative to female mice is consistent with recent data obtained in a single-choice operant paradigm in rats^20^ and with mice on a free choice assay^21^, though recent data suggests that these preferences can change in the face of stress^22^. The overall lower poke rates and longer interpoke intervals observed in female mice may suggest that females are less motivated to seek out interactions and may find social interaction less rewarding. Consistent with this idea, we found that female mice did not show robust changes in choice behavior in response to longer reward-down times. Similarly, previous work has also shown that several strains of female mice are insensitive to the effects of acute social isolation^5^. These results could indicate that the neurobiological mechanisms that influence when and how social choices are being made are less easily perturbed in female mice. On the other hand, social isolation has been shown to increase social withdrawal in female mice, while increasing aggression in male mice (Tan et al. 2021). Given that male and female mice used as subjects in this study were socially isolated throughout the experiment, it’s possible that sex differences in behavior are also influenced by differences induced by isolation.

Using normative modeling to simulate social choices(which makes predictions about how an individual ought to behave, rather than making specific predictions about implementation) allowed us to determine whether our data most closely reflected choice structure driven by multiple, parallel drives or a single social drive state. We found that the single drive model (1-Drive model) accurately predicted several key features of our data set under equal reward, female satiety, and male satiety block conditions (Fig 4). In contrast, the parallel drive model (2-Drive model) predicted that animals would be motivated to alternate choice types, and would exhibit the greatest proportion and shortest latencies for F-M and M-F (switch trials) regardless of block condition, a phenomenon that we did not observe. Importantly, no combination of parameters for the 2-Drive model was able to outperform the best fit 200 simulations of the 1-Drive model (Fig 5). These results demonstrated that in both sexes patterns of the timing and choice structure of the socially motivated choices can be explained with a generic social drive. An interesting question for future research will be to determine where the model parameters, such as choice threshold, choice bias, and interaction probability, are represented in the brain. One possibility is that midbrain dopaminergic networks may represent choice bias^20,22^, while the hypothalamic, in particular the medial preoptic area, may represent social drive^5^. While the models described here were designed for simplicity with as few parameters as possible, future work could expand these models to include other biological and physiological factors that likely influence choice behavior, such as hormone levels^23^, internal states, and the individual’s own social or experience history^24^.

Most evidence for social homeostasis in rodents has come from studies that create a negative drive state through varying lengths of isolation and explore the rebound effects^5,6,25,26^. These data echo the negative drive induced by feelings of hunger, that individuals will work to avoid^2^. In addition, recent work has shown that copulatory behavior can induce satiation in mating drives that last several days^8^. Our data expand on these findings by providing evidence that enhanced social interaction on brief timescales can act as a satiety-inducing mechanism in male mice. We observed moment-to-moment reflections of satiety in choice behavior as indicated by longer IPIs following the stated choice but not the unsated choice. demonstrating that different types of social encounters can alter satiety on varying timescales.

While midbrain and hypothalamic neurons have been implicated in these long timescale satiety mechanisms (minutes to days)^5,6,8^, it remains unclear whether they encode satiety on an interaction to interaction level. Looking for neural signatures that precede and predict social choices, predict latency to the next choice, or are modulated by the duration of social contact, may be good places to interrogate the brief satiety mechanism. Neural activity in the ventromedial hypothalamus ventrolateral area (VMHvl) has been shown to ramp up prior to social interactions, is modulated as a function of attack duration, and activation of this region can reduce interpoke-intervals in a male aggression-seeking task^14,27^, suggesting this region may be a likely candidate for driving when animals make the choice for interaction.

Overall, our data show for the first time that social interaction can act as a social satiety inducing mechanism on a moment-to-moment basis, and that social choice behavior is best represented as a generic, single-drive state. While social decision making can be complex and multifaceted, we show that social choices can be modeled with just a few, simple parameters. These results illustrate the power of using constrained, automated tasks with normative models that make predictions about self-motivated social choices to better understand social drive. The use of constrained animals in the 2-choice SOAR task prevents a feedback loop between social partners, allowing social choices to be driven by the subject animal’s behavior. At the same time, it allows for more direct social contact between the subject and stimulus mice, relative to cup-based assays or operant tasks that restrict social stimuli behind gates. Additionally, because the 2-choice SOAR task gives experimenters complete control over interaction duration, this operant assay is invaluable for answering questions related to the relationship between interaction duration and different social choices.

Finally, we list below the limitations of the current study for consideration for future experiments. While we demonstrate that male and female mice will seek out social interactions with both choices, and that longer interaction times can influence social satiety in males, we do not know the underlying motivations or affective states of the subject or stimulus animals that could influence behavior in this task. The lowering of the stimulus mice into the operant chamber is potentially stressful, which could in turn influence choice behavior. While we did not observe any overt signs of stress in the subject and reward animals, future experiments could directly test for a stress response to rule out this possibility. The male and female stimulus animals used in this study were of different strains (BALB/c and CD1, respectively). However, recent research has shown strain preference in rodents is heterogenous, with some strains showing a preference for same-strain interactions and others preferring different-strain interactions(Peng et al. 2025; Kogo et al. 2021). Furthermore, strain preference in rodents is influenced by environmental enrichment(Peng et al. 2025) and conspecific stress(Kiyokawa, Kuroda, and Takeuchi 2022). While we found that male and female CD1 mice have a preference for opposite-sex interactions, we cannot rule out the possibility that these sex differences in preference reflect differential strain preference. It was recently demonstrated that male CD1 will seek out interactions with other CD1 males (Lee et al 2024), raising the possibility that the same strain of stimulus mice could be used in future experiments. However, it is important to note that using a non-submissive strain of male reward animals may alter the rates of male-male aggression observed(Falkner et al. 2016; Minakuchi et al. 2024). Finally, while we provide evidence for a singular social drive state in the operant context, application of the model in a more naturalistic behavior setting is an important consideration for future experiments.

## MATERIALS AND METHODS

### Mice

All animal procedures were approved by the Princeton University Institutional Animal Care and Use Committee and were in accordance with National Institutes of Health standards. Animal sex was assigned based on observation of genitalia at weaning. Animals trained for the operant task were sexually naive group-housed CD1 mice between the ages of 10 and 26 weeks. Subject animals were socially isolated for at least 1 week prior to the start of training and remained isolated throughout the duration of the experiment. Male and female conspecifics used as social rewards were sexually naive, BALB/c (male) or CD1 (female) mice between the ages of 10 and 36 weeks. Mice were housed in a 12 h light-dark cycle with experiments taking place exclusively during the dark phase. Food and water were given ad libitum. All animals were bred within the Princeton University animal facilities or purchased from Taconic Biosciences.

### Video data acquisition

Operant sessions were recorded using up to three cameras (BFS-U3-13Y3m-C1.3 Mono, FLIR Systems) acquired with SpinView and controlled with a Raspberry Pi or or RX8 collected at 30 frames/sec.

### Two-choice SOAR task

The operant chamber consisted of a custom box (20” x 10”, 17”) made of acrylic boards (TAP Plastics Acrylic Sheets P95 Matte Finish - Matte Black 1/4” thick) laser cut to the specific dimensions. The two side walls were fitted with custom-made nose-poke ports, and two motorized actuators were situated behind the back wall on either side of the chamber. Two actuators (Firgelli, High Speed Linear Actuator, 12V/22 lb/18 inch), each capable of holding a reward animal, were controlled by a minicomputer (Raspberry Pi) or RX8 (tucker-davis-technologies, TDT). Nose-pokes in each port were detected by an infrared detector and emitter fitted into either side of the nose-poke-port along with a light-emitting diode (LED) at the center. On one side of the operant chamber, the left-hand port (relative to the experimental animal facing the port), activated retraction of the actuator on that side to deliver the reward animal into the operant chamber for a fixed duration of time (2-10s). On the other side of the operant chamber, poking in the right-hand port (relative to experimental animal facing the port) activated the actuator while the left-hand port was the null port. Once triggered, each social nose-poke port activated a “lock-out” period and could not be triggered again until the reward animal moved back to the original position. All nose poke and actuator movement timestamps were recorded by the Raspberry Pi or by the RX8.

Crane and tethers: Reward animals in jackets were connected to a linear actuator (Firgelli, High Speed Linear Actuator, 12V/22 lb/18 inch) via a custom arm cut from McMaster 1/4” acrylic. The mouse jacket holding a reward animal was suspended from the arm with fishing line (Piscifun Onyx Braided Fishing Line, 150lb) and magnets (KeySmart MagConnect Magnetic Key Holder) and jewelry clasps (uxcell 12mm Round Stainless Steel Spring Ring Clasps). Reward animals were constrained in custom full body jackets made from ThorLabs BK5 nylon fabric, cut on an Epilog Mini 24 30W laser. Jackets were cut with holes so that the task-engaging animal could have substantial access to the reward animal’s body surface during brief interaction intervals yet animals were comfortably cradled and supported along the length of their body during extension and retraction.

The training protocol consisted of training on a one-choice version of the task for each reward prior to the 2-choice version of the task. Experimental mice were first trained on a single side of the operant chamber, and then trained on the following side for 3-7 sessions each. During the one-choice training sessions, experimental animals were restricted to a single side of the operant chamber by an acrylic wall placed into the center of the chamber via two slots in the floor. Training on the female or male side first was counterbalanced across animals. Mice were considered “trained” and used for two-choice SOAR sessions if they reached the following criteria: a poke rate of at least 1 poke every 5 minutes, a 2:1 ratio of social to null pokes on at least one side, and performance was stable for at least two consecutive days. All mice performed a minimum of 4 two-choice SOAR sessions with equal dwell times before experiencing sessions with unequal reward durations. Different reward animals were used on each day of the experiment.

### Pose tracking and behavior analysis Behavioral annotation

To determine the types of social behaviors displayed during the 2-choice SOAR task, manual annotations of videos were conducted with Behavioral Observation Research Interactive Software (BORIS)(Friard and Gamba 2016). We annotated behavior of the subject animals, and found many trials containing bouts of sniffing directed at the anogenital region, body, or face of the stimulus mouse. We also annotated attack or fighting behavior, as defined by rapid forward movements displayed by the subject towards the reward animal during the reward-down time.

### Pose estimation

Animal pose estimation was performed with SLEAP software (v1.3.3)^12^. To create a model, mice in a subset of frames from 602 behavior videos were hand-labeled with a 15-node skeleton which labeled the following points: nose, head, left ear, right ear, trunk, tail-trunk-interface, upper segment of tail, middle-segment of tail, lower-segment of tail, and tail tip, All together, 19,594 labeled frames were split into a training and validation set using an 80:20 split. Training of the SLEAP model was done using a top-down approach with customized parameters. We then used this model to run inference on all top-view behavior videos, whereby the model produced predictions that we then hand-corrected to produce final pose estimation.**Probability of social interaction.** For each nose poke, the trial was classified by whether or not social interaction occurred. Successful social interaction was quantified by tracking the position of the subject animal’s head from moment of poke detection to the time when the reward animal was retracted. If the subject’s head entered the social zone in this time period, defined as the area immediately surrounding the reward animal, the trial was classified as successful social interaction.

### Estrous cycle staging

For both experimental and reward female mice, vaginal swabs were performed after every two-choice SOAR session to assess estrous cycle stage via cell cytology. Estrous stage was assigned from cell shape and density.

### Statistical analysis

No statistical methods were used to predetermine sample sizes, but our sample sizes are similar to those reported in previous publications. Data collection and analysis were not performed blind to the conditions of the experiments. All statistical tests for Fig 1-3 were run in Python (v3.11.7) via scipy.stats (v1.12.0) or pingouin (v0.5.4). Student’s t-test (scipy.stats.ttest_ind) or paired t-test (scipy.stats.ttest_rel) were used to assess difference in means between two groups. For determining the difference between two or more distributions, two-sample Kolmogorov-Smirnov tests (scipy.stats.k2_2samp) were conducted. Pearson’s correlation (scipy.stats.pearsonr) was used to test for linear relationship. A one-way ANOVA (pignouin.anova, pignouin.rm_anova) to assess differences in means between more than two groups, and pairwise tukey HSD test was used for post hoc tests (pignouin.pairwise_tukey). Two-way ANOVA (pignouin.mixed_anova) was used to assess differences in means between more than two groups and two independent variables. The Benjamini/Hochberg FDR method (pignouin.pairwise_comparison was used to correct for multiple comparisons (pignouin.pairwise_tests(padjust=‘FDR_BH’). See supplementary table 1 for detailed statistical parameters. For all statistical tests, significance was measured against an alpha value of 0.05. All error bars show standard error of the mean (SEM).

### Models

Simulated data was generated using a pair of simple normative models, representing a 1-Drive and a 2-Drive state and were simulated using custom scripts using Matlab R2024a. The 1-Drive model simulated motivation as a single value that can be updated by either interactions with males or females. The 2-Drive model simulates parallel motivational states for male or female that can only be updated by the choice and interaction with male or female (i.e. female motivation is updated towards satiety after interactions with females and male motivation is updated towards satiety only after interactions with males).

For both models, the motivational states of male and female subjects were modeled over a specified number of timesteps. We used 10k timesteps for figure graphic and 100k timesteps for quantification and model fits. Values of this motivational state were negative, with more negative values representing greater satiety. All motivation levels were initialized at the first timestep with random negative values for both male and female subjects. These initial values were drawn from a uniform distribution multiplied by −2, creating initial states that represent a low motivation (sated) level.

For each subsequent timestep, the motivation levels were updated using an exponential decay function. The decay rates were governed by predefined time constants, reflecting an natural increase in motivation (less satiety) over time. A single time constant was used for the 1-Drive model, while in the 2-Drive model, these values could be set independently for male and female motivation.

At each timestep, if the current motivation level exceeds a certain threshold (a single value for the 1-Drive model, but 2 separate values for male and female thresholds), the model automatically makes a “choice” for male or female. In the 1-Drive model, the choice (male vs. female) is made by comparing a randomly generated number between 0 and 1 to a “choice bias” term. If the random number is higher than the choice bias term, a choice to female is made. A choice bias set to 0.5 will result in random, but equal choices to male and female rewards. For the 2-Drive model, the choice is determined by which motivational state (male or female) is the one to reach its threshold. Since individuals do not always interact after making the choice, we also determine the likelihood of interaction after each choice by comparing a random number drawn between 0 and 1 to an “interaction probability” value. For both 1-Drive and 2-Drive models, the interaction probability can be set independently for male and female choices.

To differentially model the effects of long and short dwell periods (reward-down time), the amount of input given to the model after a choice is varied. For each threshold crossing, motivation was reduced by a randomly determined value within a specified range representing the “dwell time” effect. Long and short dwell times were specified by a range of values. Large inputs to the model (representing a larger deviation towards satiety and long reward-down times, were sampled from within the range of 2-2.5, while small inputs to the model (representing a smaller deviation towards satiety and short reward-down time) were sampled from the range of 0.4-0.5. If no interaction occurred, the motivation was reduced by a fixed value representing the lack of interaction and input given to the model was set to 0.1.

Throughout the simulation, instances where the motivation crossed the threshold and triggered an event were recorded. These events were logged in separate vectors for male and female subjects. Additionally, the total counts of these events were recorded for each sex.

The latency of each event was calculated and sorted in ascending order. The order of these events was used to determine the sequence of interactions between male and female subjects. Transition probabilities between successive events were calculated, categorized into four types: female-to-female (FF), female-to-male (FM), male-to-male (MM), and male-to-female (MF). These transition probabilities were expressed as proportions of the total number of transitions.

Finally, the average latencies between successive events for each transition type were computed, providing insights into the temporal dynamics of interaction sequences in the simulated population. These latencies were averaged across all occurrences of each transition type to generate summary statistics for the timing of these interactions.

We generated sample data from the 1-Drive and 2-Drive model using a set of parameters that produce data that was visually similar to the behavioral data. For the 1-Drive model, we used a time constant of 50, probability of interaction with male and female of 0.7, and a threshold of −0.01, and a choice bias of 0.5 or 0.6. For the 2-Drive model, we similarly used time constants for male and female motivation of 50, probability of interaction with male and female of 0.7, and a threshold of −0.01 for both male motivation and female motivation.

To iteratively screen combinations of parameters for both 1-Drive and 2-Drive models and compare the simulated data to behavioral data, we systematically simulated data with a range or parameter values and regressed the simulated vector of choice probabilities and normalized latencies with behavior data from each satiety manipulation. For behavioral data, latency vectors were normalized using the max latency value across all three satiety manipulations (equal, male long and female long).

## Code Availability

Code for model simulations is available on GitHub repository:

https://github.com/FalknerLab/2SOAR. Behavioral and pose data available upon reasonable request

## Acknowledgments

We thank members of the Falkner laboratories for useful discussions, Gloria Liu for graphics, G, Seiler, L. Sirrs and A. Le for help with mice. Funding was from DP2MH126375 (to A.L.F.), NIH R01MH126035 (to A.L.F.), NIH F32MH136673 (to N.R.M), NYSCF (to A.L.F.), SCGB (to A.L.F.), Klingenstein Foundation (to A.L.F.), Nakajima Foundation (T.M.).

A.L.F. is a New York Stem Cell Foundation Robertson Investigator.

---

**Supplemental Figure 1.**
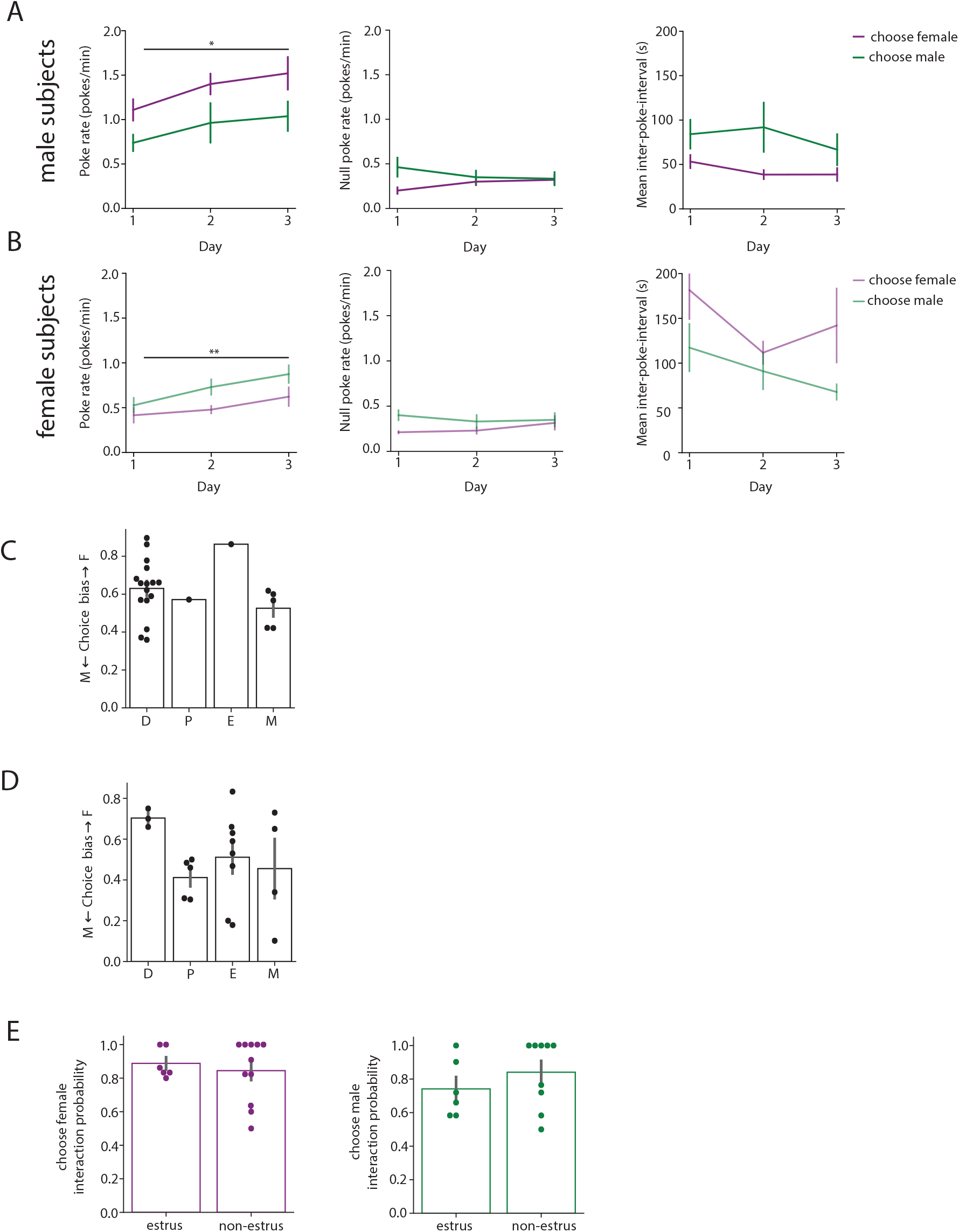
**(A)** Average poke rate (left), null poke rate (middle) and mean interpoke-interval (right) for female choices (purple) and male choices (green) across the first three days of the 2-choice SOAR task in male subjects. (Two-way mixed ANOVA, pairwise comparison with FDR correction). **(B)** Same as **(A)** but for female subjects. **(C)** Influence of reward female estrous state on male subject choice bias (Kruskal-Wallis test, *p* = 0.276). **(D)** Influence of female subject estrous state on choice bias (Kruskal-Wallis test, *p* = 0.207). **(E)** There is no influence of estrus state on interaction probability following female choices (left, *t(15) = 0.531, p = 0.603*) or male choices (*right, t*(13) = 0.981, *p* = 0.344). Plots in A-B are presented as mean ± s.e.m. * p < 0.05, *\*\** p < 0.01.

**Supplementarl Figure 2.**
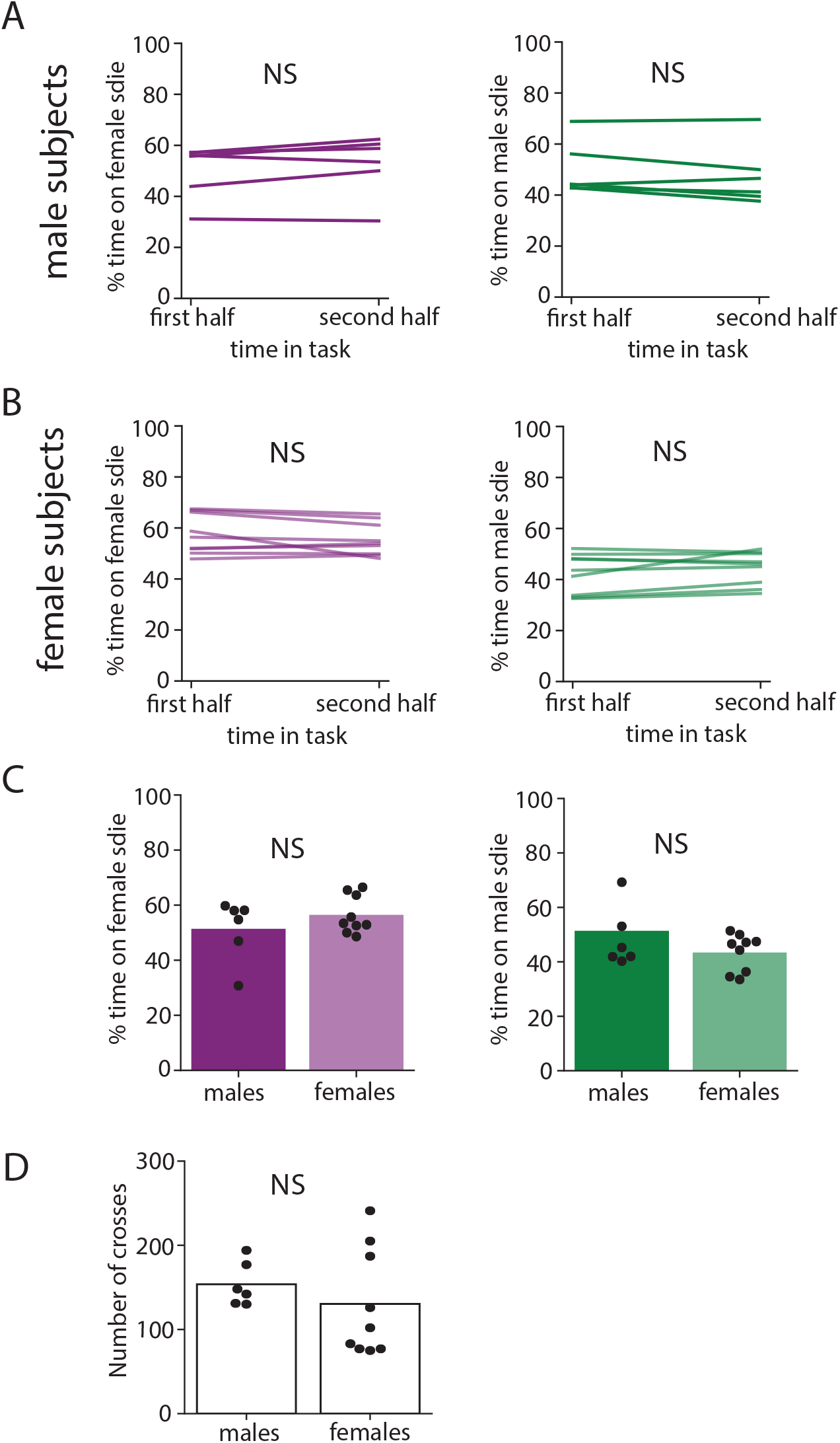
**(A)** Percent time spent on the female side (left; *p* = 0.158) or male side (right; *p* = 0.159) of the operant chamber during the first and second half of the task in male subjects. Independent samples t-test. **(B)** Same as **A** but for female subjects (repeated measures t-test; female side: *p* = 0.173; male side: *p* = 0.174). **(C)** Comparison of time spent on the female side (left, *p* = 0.285) or male side (right, *p* = 0.285) of the operant chamber between males and females. **(D)** Number of crosses in the operant chamber between male and females (independent samples t-test, *p* = 0.417). NS, nonsignificant.

**Supplemental Video 1.**

Example tracked video from male mouse performing sequential choices in 2-choice SOAR task.

**Supplementary Table 1.**

Information outlined in each section contains the figure number (second column), figure description (third column), statistical test used (fourth column), test outcome (fifth column), and posthoc test (sixth column). Final reported p-values are in the “Outcome” or “Posthoc test” column.

